# Competing neural representations of choice shape evidence accumulation in humans

**DOI:** 10.1101/2022.10.03.510668

**Authors:** Krista Bond, Javier Rasero, Raghav Madan, Jyotika Bahuguna, Jonathan Rubin, Timothy Verstynen

## Abstract

Changing your mind requires shifting the way streams of information lead to a decision. Using *in silico* experiments we show how the cortico-basal ganglia-thalamic (CBGT) circuits can feasibly implement shifts in the evidence accumulation process. When action contingencies change, dopaminergic plasticity redirects the balance of power, both within and between action representations, to divert the flow of evidence from one option to another. This finding predicts that when competition between action representations is highest, the rate of evidence accumulation is lowest. We then validate this prediction in a sample of *homo sapiens* as they perform an adaptive decision-making task while whole-brain hemodynamic responses are recorded. These results paint a holistic picture of how CBGT circuits manage and adapt the evidence accumulation process in mammals.

**One-sentence Summary:** Interactions between cortical and subcortical circuits in the mammalian brain flexibly control the flow of information streams that drive decisions by shifting the balance of power both within and between action representations.

## Introduction

Choice is fundamentally driven by information. The process of deciding between available actions is continually updated using incoming sensory signals, processed at a given accumulation rate, until sufficient evidence is reached to trigger one action over the other (*1, 2*). The parameters of this evidence accumulation process are highly plastic, adjusting to both the reliability of sensory signals (*3–7*) and previous choice history (*8–13*), to balance the speed of a given decision with local demands to choose the right action.

We recently showed how environmental change influences the decision process by periodically switching the reward associated with a given action in a 2-choice task (*7*). This reward contingency change induces competition between old and new action values, forcing a change-of-mind about the most rewarding option. This internal competition prompts humans to dynamically reduce the rate at which they accumulate evidence (drift-rate, *v*, in a normative drift diffusion model, DDM (*2*)) and sometimes also increases the threshold of evidence they need to trigger an action (boundary height, *a*) (*7*). The result is a change of the decision policy to a slow, exploratory state. Over time feedback-learning pushes the system back into an exploitative state until the environment changes again (see also (*11*)) and (*12*)).

Here we investigate the underlying neural mechanisms that drive dynamic decision policies in a changing environment. We start with a set of theoretical experiments, using biologically realistic spiking network models, to test how the cortico-basal ganglia-thalamic (CBGT) circuits influence the evidence accumulation process (*14–18*). These experiments both explain previous results (*7*) and make specific predictions as to how competition between action representations drives changes in the decision policy. We then test these predictions in humans using a high-powered, within-participant neuroimaging design, collecting data over thousands of trials where action-outcome contingencies change on a semi-random basis.

## Results

### CBGT circuits can control decision parameters under uncertainty

Both theoretical (*9, 12, 14, 19–21*) and experimental (*18*) evidence suggest that the CBGT circuits play a critical role in the evidence accumulation process (for a review see (*22*)). The canonical CBGT circuit (Fig. 1A) includes two dissociable control pathways: the direct (facilitation) and indirect (suppression) pathways (*23, 24*). A critical assumption of the canonical model is that the basal ganglia are organized into multiple “channels” mapped to specific action representations (*25,26*), each containing a direct and indirect pathway. While a strict, segregated action channel organization may not accurately reflect the true underlying circuitry, striatal neurons have been shown to organize into task-specific spatiotemporal assemblies that qualitatively reflect independent action representations (*27–31*). Within these action channels, activation of the direct pathway, via cortical excitation of D1-expressing spiny projection neurons (SPNs) in the striatum, releases GABAergic signals that can suppress activity in the CBGT output nucleus (internal segment of the globus pallidus, GPi, in primates or substantia nigra pars reticulata, SNr, in rodents) (*26, 32–34*). This relieves the thalamus from tonic inhibition, thereby exciting postsynaptic cortical cells and facilitating action execution. Conversely, activation of the indirect pathway via D2-expressing SPNs in the striatum controls firing in the external segment of the globus pallidus (GPe) and the subthalamic nucleus (STN), resulting in strengthened basal ganglia inhibition of the thalamus. This weakens drive to postsynaptic cortical cells and reduces the likelihood that an action is selected in cortex.

**Fig. 1.**
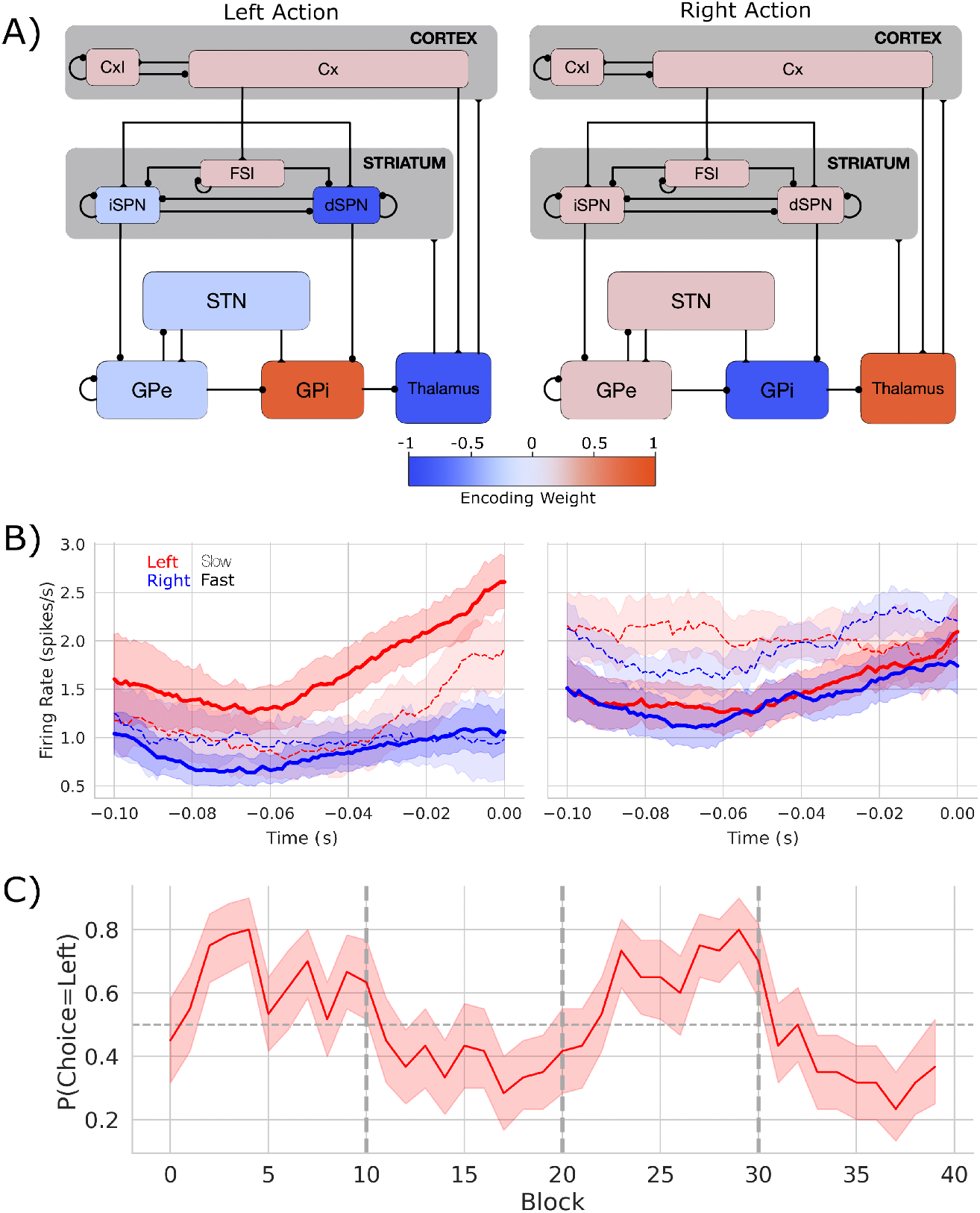
Biologically based CBGT network dynamics and behavior. A) Each CBGT nucleus is organized into left and right action channels with the exception of a common population of striatal fast spiking interneurons (FSIs) and another of cortical interneurons (CxI). Values show encoded weights for a left action. Network schematic adapted from (*19*). B) Firing rate profiles for D1-SPNs (left panel) and D2-SPNs (right panel) prior to stimulus onset (t=0) for a left choice. D1-SPN activity in left and right channelsis shown in red and blue, respectively. Thick solid lines represent fast trials (short RTs) and thin dashed lines represent slow trials (long RTs). C) Choice probability for the CBGT network model. The reward for left and right actions changed every 10 trials, marked by vertical dashed lines. The horizontal dashed line represents chance performance.

Critically, the direct and indirect pathways converge in the GPi/SNr (*35, 36*). This suggests that these pathways compete to control whether each specific action is selected (*37*). The apparent winner-take-all selection policy and action-channel like coding (*27–31*) also imply that action representations themselves compete. Altogether, this neuroanatomical evidence suggests that competition both between and within CBGT pathways controls the rate of evidence accumulation during decision making (*12, 15, 19*).

To illustrate this process, we designed a spiking neural network model of the CBGT circuits, shown in Fig. 1A, with dopamine-dependent plasticity occurring at the corticostriatal synapses (*17, 38*). The network performed a probabilistic 2-arm bandit task with switching reward contingencies ((*7*); see Supp. Methods). The experimental task followed the same general structure as our prior work (*7*). In brief, the network selected one of two targets, each of which returned a reward according to a specific probability distribution. The relative reward probabilities for each target were held constant at 75% and 25% and the action-outcome contingency was changed every 10 trials, on average. For the purpose of this study we focus primarily on the neural and behavioral effects that occur around the switching of the optimal target. We used four different network instances (see Supp. Methods) as a proxy for simulating individual differences over human participants.

Figure B shows the firing rates of dSPNs and iSPNs in the left action channel, time-locked to selection onset (when thalamic units exceed 30Hz, t=0), for both fast (*<*196ms) and slow (*>* 314.5ms) decisions. As expected, the dSPNs show a ramping of activity as decision onset is approached and the slope of this ramp scales with response speed. In contrast, we see that iSPN firing is sustained during slow movements and weakly ramps during fast movements. However, iSPN firing was relatively insensitive to left versus right decisions. This is consistent with our previous work showing that differences in direct pathways track primarily with choice while indirect pathway activity modulates overall response speeds (*12, 19*) as supported by experimental studies (*39–41*).

We then modeled the behavior of the CBGT network using a hierarchical version of the DDM (*42*), a canonical formalism for the process of evidence accumulation during decision-making (*2*) (Fig. 2A). This model returns four key parameters with distinct influences on evidence accumulation. The drift rate (*v*) represents the rate of evidence accumulation, the boundary height (*a*) represents the amount of evidence required to cross the decision threshold, nondecision time (*t*) is the delay in the onset of the accumulation process, and starting bias (*z*) is a bias to begin accumulating evidence for one choice over another (see Methods section).

**Fig. 2.**
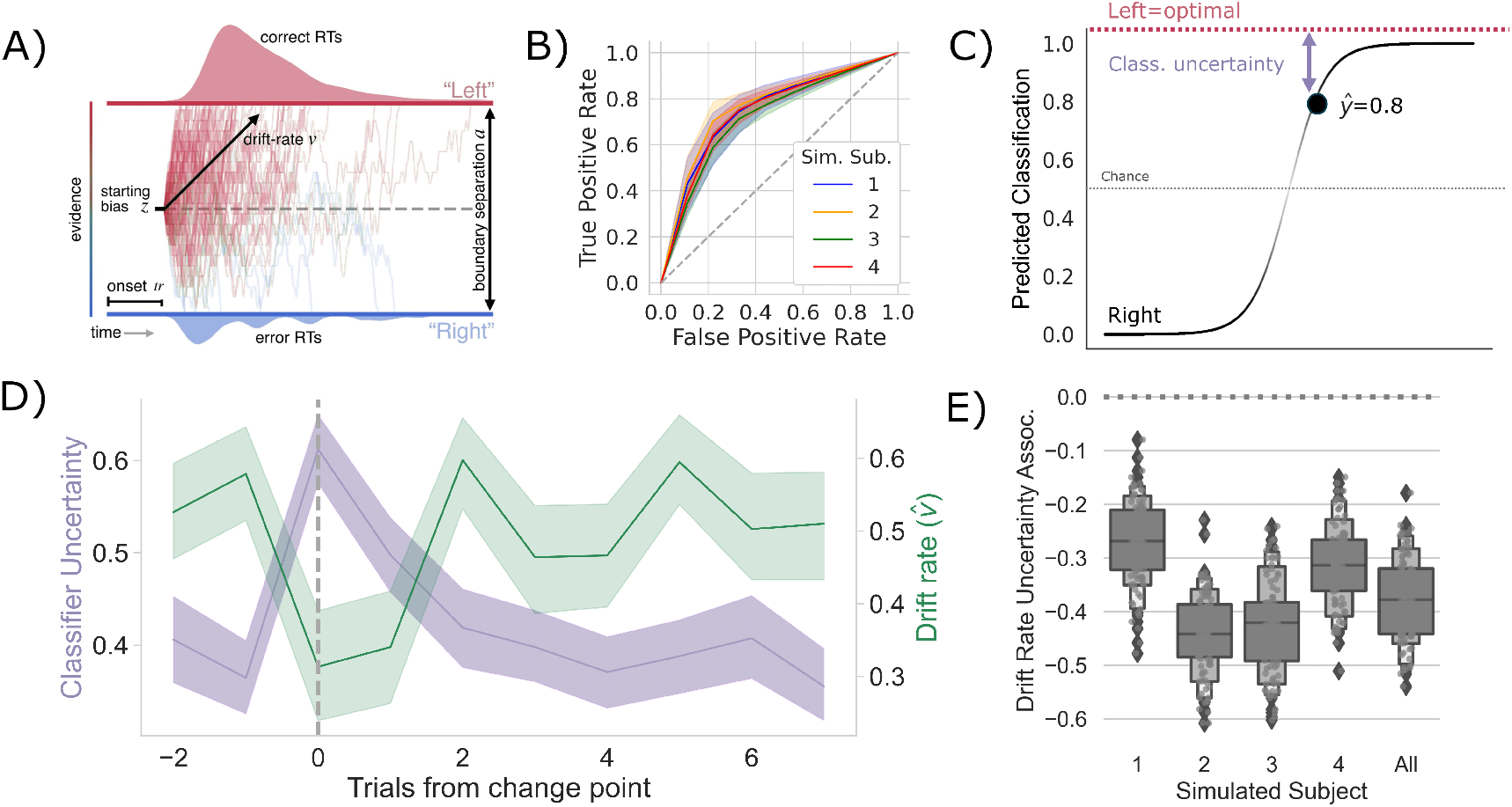
Competition between action plans *should* drive evidence accumulation. A) Decision parameters were estimated by modeling the joint distribution of reaction times and responses within a drift diffusion framework. B) Classification performance for single-trial left and right actions shown as an ROC curve. The gray dashed line represents chance performance. C) Predicted left and right responses. The distance of the predicted response from the optimal choice represents classifier uncertainty for each trial. For example, here the predicted probability of a left response on the first trial 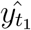 is 0.8. The distance from the optimal choice on this trial and, thereby, the classifier uncertainty 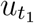, is 0.2. D) Change-point-evoked classifier uncertainty (lavender) and drift rate (green). The change point is marked by a dashed line. E) The association between classifier uncertainty and drift rate. Results for individual participants are presented along with aggregated results.

We tracked internal estimates of action-value and environmental change using trial-by-trial estimates of two ideal observer parameters, the belief in the value of the optimal choice (Δ*B*) and change point probability (Ω), respectively (see (*3, 7*) and Methods for details). Using these estimates, we evaluated how a suspected change in the environment and the belief in optimal choice value influenced underlying decision parameters. Consistent with prior observations in humans (*7*) we found that both *v* and *a* were the most pliable parameters across experimental conditions for the network. Specifically, we found that the model mapping Δ*B* to drift rate and Ω to boundary height and the model mapping Δ*B* to drift rate provided equivocal best fits to the data over human participants (Δ*DIC*_null_ = −29.85 ± 12.76 and Δ*DIC*_null_ = −22.60 ± 7.28, respectively; see (*43*) and Methods for guidelines on model fit interpretation). All other models failed to provide a better fit than the null model (Supp. Table 2). Consistent with prior work (*7*), we found that the relationship between Ω and the boundary height was unreliable (mean *β*_*a*∼Ω_ = 0.069±0.152; mean *p* = 0.232±0.366). However, drift rate reliably increased with Δ*B* in three of four participants (mean *β*_*v*∼Δ*B*_ = 0.934 ± 0.386; mean *p <* 0.001; 4*/*4 participants *p <* 0.001; Supp. Table 3).

These effects reflect a stereotyped trajectory around a change point, whereby *v* immediately plummets and *a* briefly increases, with *a* quickly recovering and *v* slowly growing as reward feedback reinforces the new optimal target (*7*). Because prior work has shown that the change in *v* is more reliable than changes in *a* (*7*) and because *v* determines the direction of choice, we focus the remainder of our analysis on the control of *v*.

To test whether these shifts in *v* are driven by competition within and between action channels, we predicted the network’s decision on each trial using a LASSO-PCR classifier trained on the pre-decision firing rates of the network (see Measuring neural action representations). The cross-validated accuracy for the four simulated participants is shown in Figure B. This model was able to predict the chosen action with ≈ 70% accuracy (72-77%) for each simulated participant, with an overall accuracy of ≈ 74%. Examining the encoding pattern in the simulated network, we see lateralized activation over left and right action channels (Fig. 1A), with opposing weights in GPi and thalamus, and, to a lesser degree, contralateral encoding in STN and in both indirect and direct SPNs in striatum. We do not observe contralateral encoding in cortex, which likely reflects the emphasis on basal ganglia structures and lumped representation of cortex in the model design.

To quantify the competition between action channels, we took the unthresholded prediction from the LASSO-PCR classifier, *ŷ*_*t*_, and calculated its distance from the optimal target (i.e., target with the highest reward probability) on each trial (Supp. Fig. 3; Fig. 2C). This provided an estimate of the classifier’s uncertainty driven by the separability of pre-decision activity across action channels. In other words, the distance from the optimal target should increase with increased co-activation of circuits that represent opposing actions.

**Fig. 3.**
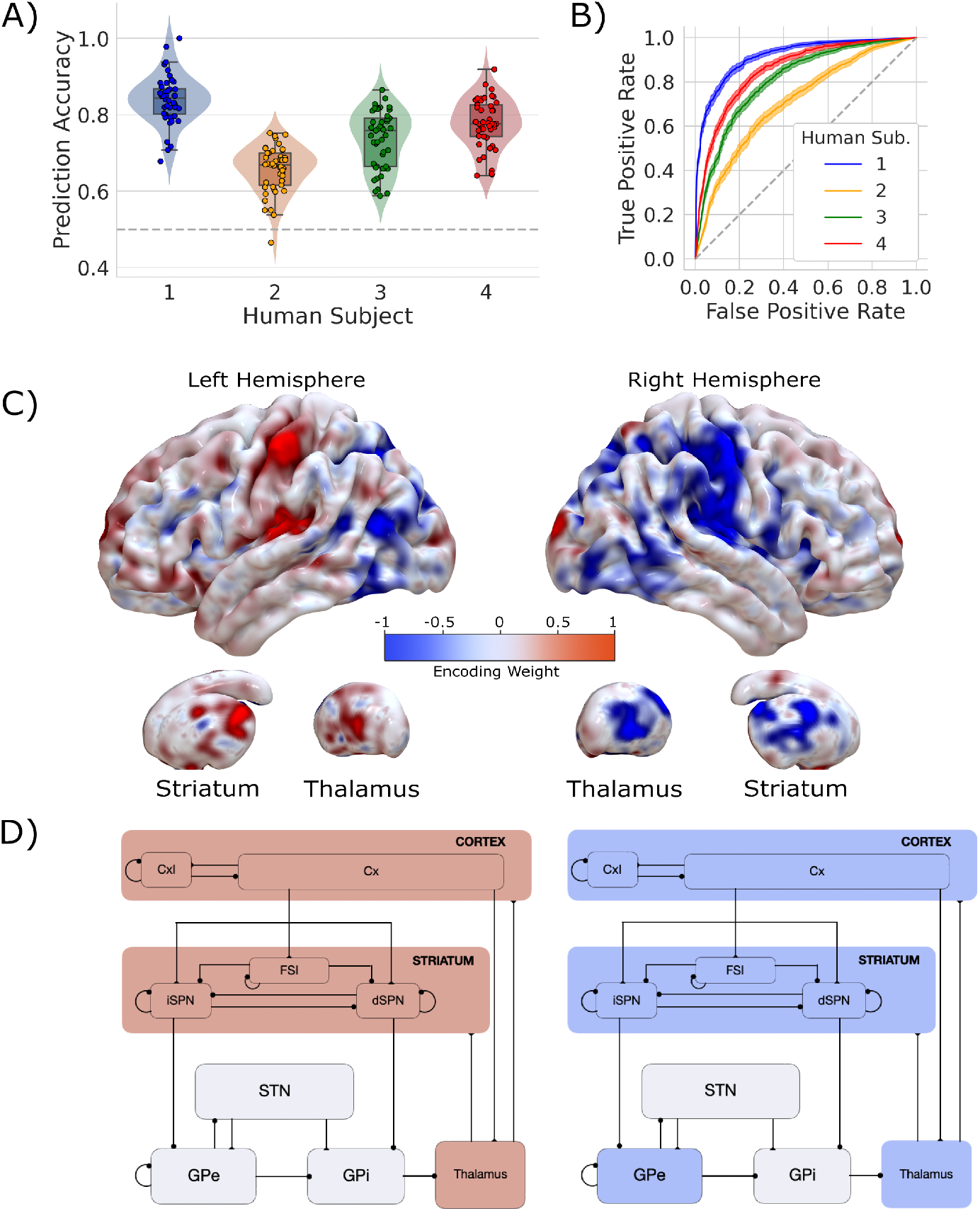
Single-trial prediction of action plan competition in humans. A) Overall classification accuracy for single-trial actions for each participant. Each point corresponds to the performance for each cross-validation fold. B) Classification performance for single-trial actions shown as an ROC curve. The gray dashed line represents chance performance. C) Participant-averaged encoding weight maps in standard space for both hemispheres. D) The mean encoding weights within each CGBT node in both hemispheres. See encoding weight scale above for reference.

If the competition in action channels is also driving *v*, then there should be a negative correlation between the classifier’s uncertainty and *v*, particularly around a change point. Indeed, this is exactly what we see (Fig. 2D). In fact, the classifier’s uncertainty and *v* are consistently negatively correlated across all trials in every simulated participant and in aggregate (Fig. 2E). Thus, in our model of the CBGT pathways, competition between action representations drives changes in *v* in response to environmental change.

### *Homo sapiens* adapt decision policies in response to change

To test the predictions of our model, a sample of primates (*Homo sapiens*, n=4) played a dynamic two-armed bandit task under experimental conditions similar to those used for the simulated CBGT network and prior behavioral work (*7*) as whole brain hemophysiological signals were recorded using functional magnetic resonance imaging (fMRI). On each trial, participants were presented with a male and female Greeble (*44*). The goal was to select the Greeble most likely to give a reward. Selections were made by pressing a button with their left or right hand to indicate the left or right Greeble on the screen. We collected 2700 trials over 45 runs from nine separate imaging sessions per participant. Consistent with our within-participant design, statistical analyses estimated effects on a single-participant basis.

Overall, speed and accuracy across conditions matched what we observed in previous experiments (Experiment 2 in (*7*)). Specifically, we see a consistent effect of change point on both RT and accuracy that matches the behavior of our network (Supp. Fig. 2; Supp. Table 1).

To address how a change in the environment shifted underlying decision dynamics, we used a hierarchical DDM modeling approach (*42*) as we did with the network behavior (see Methods for details). Given previous empirical work (*7*) and the results from our CBGT network model showing that only *v* and, less reliably, *a* respond to a shift in the environment (*7*), we focused our subsequent analysis on these two parameters. Consistent with the predictions from our CBGT model, we found equivocal fits for the model mapping both Δ*B* to *v* and Ω to *a* and a simpler model mapping Δ*B* to *v* (see Supp. Table 2 for average results). This pattern was fairly consistent at the participant level, with 3/4 participants showing Δ*B* modulating *v* (Supp. Table 3). These results suggest that as the belief in the value of the optimal choice approaches the reward value for the optimal choice, the rate of evidence accumulation increases.

Taken altogether, we confirm that humans rapidly shift how quickly they accumulate evidence (and, to some degree, how much evidence they need to make a decision) in response to a change in action-outcome contingencies. This mirrors the decision parameter dynamics predicted by the CBGT model. We next evaluated how this change in decision policy tracks with competition in neural action representations.

### Measuring action representations in the brain

To measure competition in action representations, we first needed to determine how individual regions (i.e., voxels) contribute to single decisions. For each participant, trial-wise responses at every voxel were estimated by means of a general linear model (GLM), with trial modeled as a separate condition in the design matrix. Therefore, the 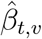 estimated at voxel *v* reflected the magnitude of the evoked response on trial *t*. As in the CBGT model analysis, these whole-brain, single-trial responses were then submitted to a LASSO-PCR classifier to predict left/right response choices. The performance of the classifier for each participant was evaluated with a 45-fold cross-validation, iterating through all runs so that each one corresponded to the hold-out test set for one fold.

Our classifier was able to predict single trial responses well above chance for each of the four participants (Fig. A and B), with mean prediction accuracy ranging from 65% to 83% (AUCs from 0.72 to 0.92). Thus, as with the CBGT network model, we were able to reliably predict trial-wise responses for each participant. Fig 3C shows the average encoding map for our model as an illustration of the influence of each voxel on our model predictions (Supp. Fig. 4 displays individual participant maps). These maps effectively show voxel-tuning towards rightward (blue) or leftward (red) responses. Qualitatively, we see that cortex, striatum, and thalamus all exhibit strongly lateralized influences on contralateral response prediction. Indeed, when we average the encoding weights in terms of principal CBGT nuclei (Fig. 3D), we confirm that these three regions largely predict contralateral responses. Supp. Fig. 5 provides a more detailed summary of the encoding weights across multiple cortical and subcortical regions.

**Fig. 4.**
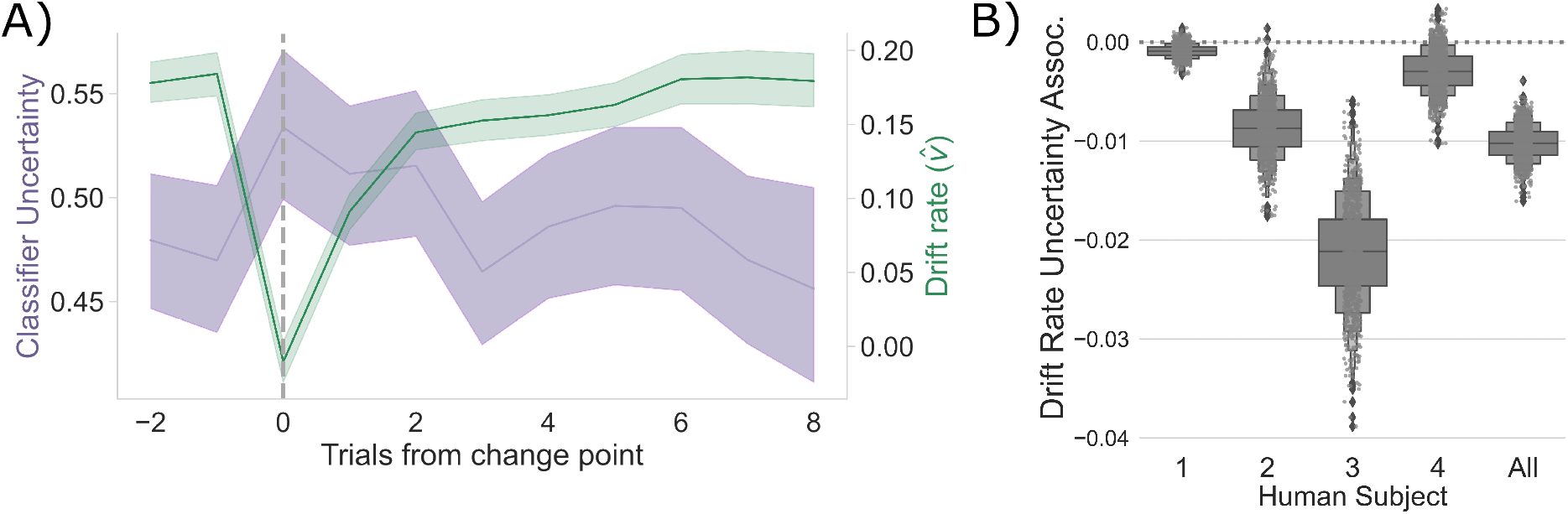
Competition between action plans drives evidence accumulation in humans. A) Classifier uncertainty (lavender) and estimated drift rate (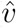; green) dynamics. B) The association between classifier uncertainty and drift rate by participant and in aggregate.

These results show that we can reliably predict single-trial choices from whole-brain hemodynamic responses for individual participants. Further, key regions of the CBGT pathway contribute to these predictions. Next, we set out to determine whether competition between these representations for left and right actions correlates with changes in the drift rate, as predicted by the CBGT network model (Fig. 2C).

### Competition between action representations may drive drift-rate

To evaluate whether competition between action channels correlates with the magnitude of *v* on each trial, as the CBGT network predicts (Fig. 2C), we focused our analysis on trials surrounding the change point, following analytical methods identical to those described in the previous section and shown in Fig. 2C.

Consistent with the CBGT network model predictions, following a change point, *v* shows a stereotyped drop and recovery as observed in the CBGT network (Fig. 2C) and prior behavioral work (*7*) (Fig. 4A). This drop in *v* tracked with a relative increase in classifier uncertainty, and subsequent recovery, in response to a change in action-outcome contingencies (mean bootstrapped *β*: −0.021 to −0.001; *t* range: −3.996 to −1.326; *p*_*S*1_ = 0.057, *p*_*S*2_ *<* 0.001; *p*_*S*3_ *<* 0.001; *p*_*S*3_ = 0.080, *p*_All_ *<* 0.001). As with the CBGT network simulations (Fig. 2D), we also observe a consistent negative correlation between *v* and classifier uncertainty over all trials, irrespective of their position to a change point, in each participant and in aggregate (Fig. 4B; Spearman’s *ρ* range: −0.08 to −0.04; *p* range: *<* 0.001 to 0.043).

These results clearly suggest that, as predicted by our CBGT network simulations and prior work (*12, 17, 45*), competition between action representations drives changes in the rate of evidence accumulation during decision making in humans.

## Discussion

We investigated the underlying mechanisms that drive shifts in decision policies when the rules of the environment change. We first tested an implementation-level theory of how CBGT networks contribute to changes in decision policy parameters. This theory predicted that the rate of evidence accumulation is driven by competition across action representations. Using a high-powered within-participants fMRI design conducted with four human primates, wherein each participant served as an independent replication test, we found evidence consistent with our CBGT network simulations. Specifically, as action-outcome contingencies change, thereby increasing uncertainty of optimal choice, decision policies shift with a rapid decrease in the rate of evidence accumulation, followed by a gradual recovery to baseline rates as new contingencies are learned (see also (*7*)). These results empirically validate prior theoretical and computational work predicting that competition between neural populations encoding distinct actions modulates how information is used to drive a decision (*9, 12, 14, 20, 21*).

Our findings here align with prior work on the role of competition in the regulation of evidence accumulation. In the decision-making context, the ratio of dSPN to iSPN activation *within* an action channel has been linked to the drift-rate of single-action decisions (*14–16, 37*). In the motor control context, this competition manifests as movement vigor (*46–48*). Yet, our results show how competition *across* channels drives drift-rate dynamics. So how do we reconcile these two effects? Mechanistically, the strength of each action channel is defined by the relative difference between dSPN and iSPN influence. In this way, competition across action channels is defined by the relative balance of direct and indirect pathway activation within each channel. Greater direct vs. indirect pathway competition in one action channel, relative to another, makes that action decision relatively slow and reduces the overall likelihood that it is selected. This mechanism is consistent with prior theoretical (*12, 45*) and empirical work (*18*).

While our current work postulates a mechanism by which changes in action-outcome contingencies drive changes in evidence accumulation through plasticity within the CBGT circuits, the results presented here are far from conclusive. For example, our model of the underlying neural dynamics predicts that the certainty of individual action representations is encoded by the competition between direct and indirect pathways (see also (*12, 38, 45*)). Thus, external perturbation of dSPN (or iSPN) firing, say with optogenetic methods, during decision-making should causally impact the evidence accumulation rate and, subsequently, the speed (or slow) the speed at which the new action-outcome contingencies are learned. Indeed, there is already some evidence for this outcome (see (*18*), but also (*49*) for contrastive evidence).

Our model, however, has very specific predictions with regards to disruptions of each pathway within an action representation. Disrupting the *balance* of dSPN and iSPN efficacy should selectively impact the drift-rate (and, to a degree, onset bias; see (*45*)), while non-specific disruption of global iSPN efficacy across action representations should selectively disrupt boundary height (and, to a degree, accumulation onset time; see again (*45*)).

Thus, increasing the difference between dSPN and iSPN firing in the channel representing the new optimal-action, say by selective excitation of the relevant dSPNs, should speed up the time to resolve the credit assignment problem during learning. This would result in faster and more accurate learning following an environmental change and lead to characteristic signatures in the distribution of reaction times, as well as choice probabilities, reflective of a shift in evidence accumulation rate. Of course, testing these predictions is left to future work.

## Conclusion

As the world changes and certain actions become less optimal, successful behavioral adaptation requires flexibly changing how sensory evidence drives decisions. Our simulations and hemophysiological experiments in human primates show how this process can occur within the CBGT circuits. Here, a shift in action-outcome contingencies induces competition between encoded action plans by modifying the relative balance of direct and indirect pathway activity in CBGT circuits,both within and between action channels, slowing the rate of evidence accumulation to promote adaptive exploration. If the environment subsequently remains stable, then this learning process accelerates the rate of evidence accumulation for the optimal decision by increasing the strength of action representations for the new optimal choice. This highlights how these macroscopic systems promote flexible, effective decision-making under dynamic environmental conditions.

## Acknowledgements

We thank all members of the CoAx Lab and collaborators for their feedback on the development of this work.

## Author Contributions

K.B.: Conceptualization, Data curation, Formal analysis, Investigation, Methodology, Project administration, Software, Visualization, Writing - original draft, Writing - review and editing;

JR: Formal analysis, Software, Visualization, Writing - original draft, Writing - review and editing;

RM: Data curation, Formal analysis, Software, Writing - review and editing;

JB: Formal analysis, Software, Writing - review and editing;

JR: Conceptualization, Writing - review and editing;

TV: Conceptualization, Formal analysis, Funding acquisition, Investigation, Project administration, Resources, Supervision, Validation, Writing - review and editing

## Data and Materials Availability

Behavioral data and computational derivatives are publically available here. Raw and preprocessed hemodynamic data, in addition to physiological measurements collected for quality control, are available here.

## Supplementary Materials

Materials and Methods Supplementary Text

Figs. S1 to S5

Tables S1 to S8

Supp. References (0)

No supplementary data included (see Data Availability section above)

### Figures

**Supp. Fig. 1.**
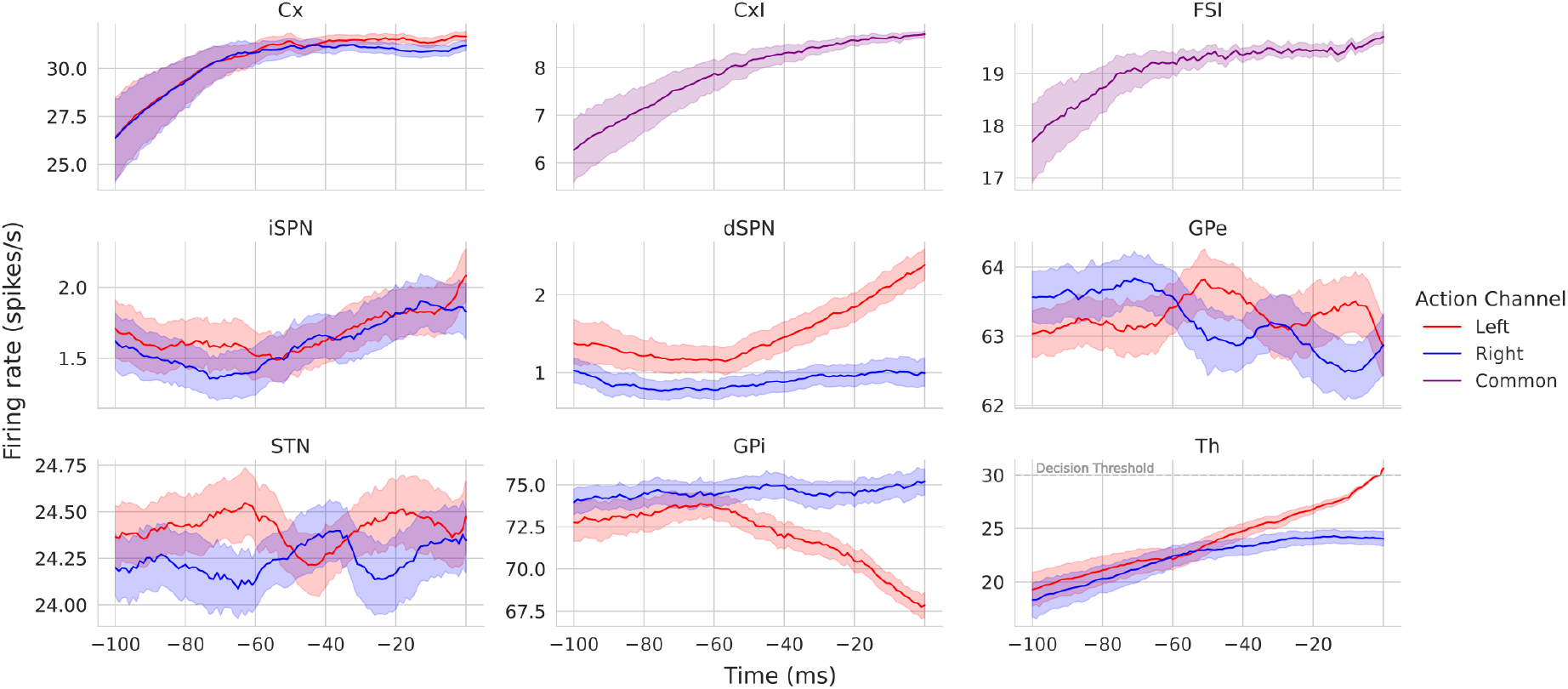
Simulated CBGT nuclei firing rates for a left decision. Each panel shows the firing rates for each CBGT nucleus 100 ms prior to a left decision. The decision threshold for thalamus (30 spikes/second) is marked with a horizontal gray line. Note that the y axes have different limits for each nucleus due to differences of scale in their firing rates.

**Supp. Fig. 2.**
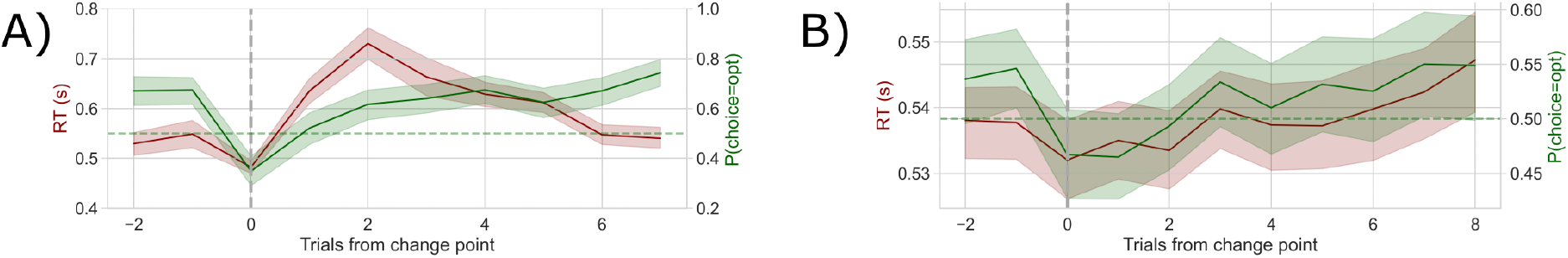
Simulated and human behavior. Change point evoked reaction times are shown in red and accuracy, or the probability of selecting the optimally rewarding choice, is shown in green. Chance is marked as a green horizontal dashed line. The change point is marked by the vertical gray line. A) Simulated behavior. B) Human behavior.

**Supp. Fig. 3.**
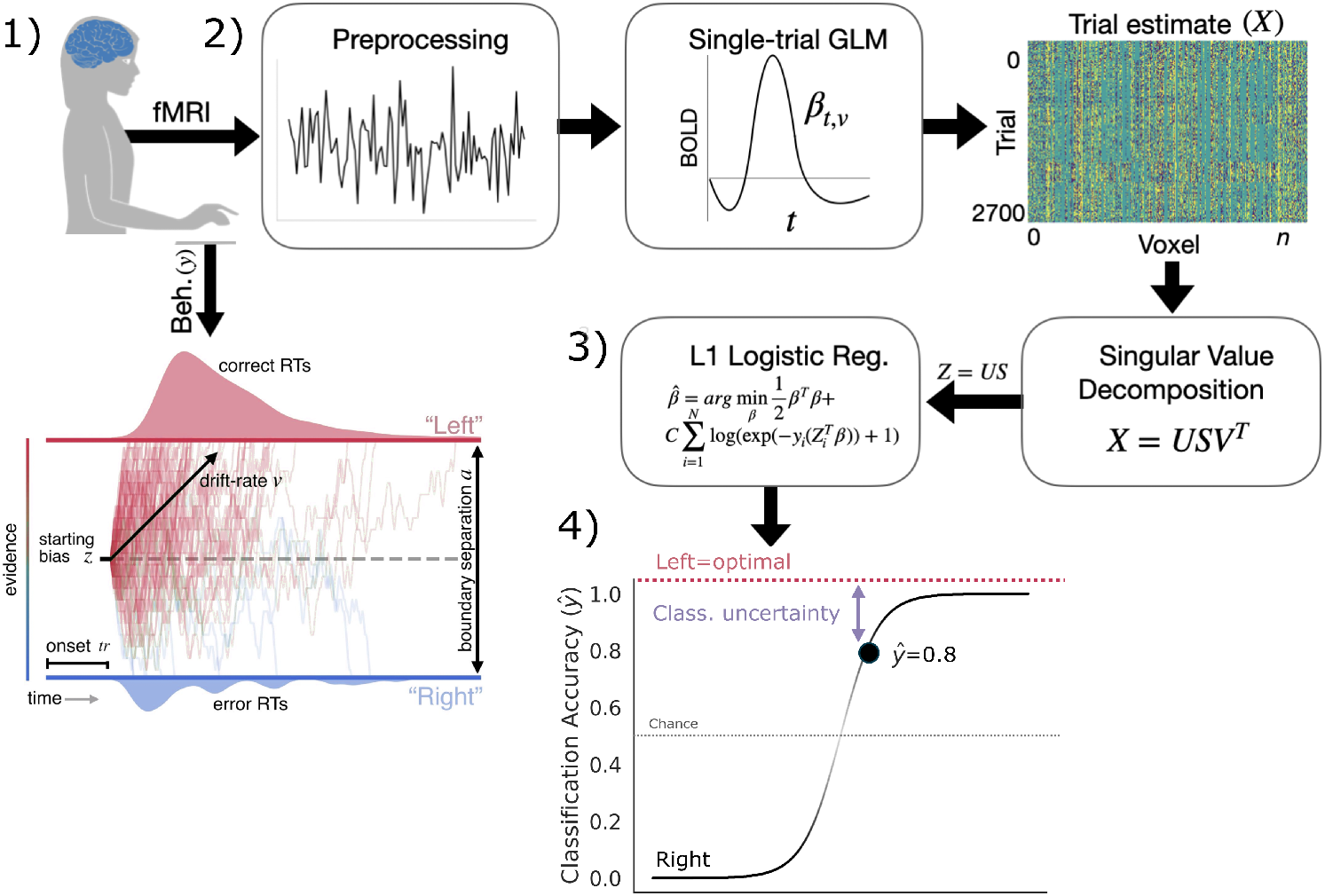
Analysis method. Step 1. Behavioral response collection and DDM (Drift Diffusion Model) parameter estimation. In the case of the simulated CBGT network, this step involved simulating responses to experimental manipulations. Step 2. Preprocessing and single-trial estimates of the hemodynamic response. Step 3. Singular Value Decomposition and Logistic regression with an L1 penalty. After crossvalidation, this outputs a predicted response (left or right), here coded as 0 or 1. Step 4. Calculating classifier uncertainty from cross-validated response prediction. The further the predicted response from the inflection point of the logistic function, the more certain the prediction. The distance of this predicted response from the optimal choice represents classifier uncertainty for each trial. Here, the predicted probability of a left response 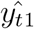 is 0.2. The distance from the optimal choice on this trial, and, thereby, the classifier uncertainty is 0.2.

**Supp. Fig. 4.**
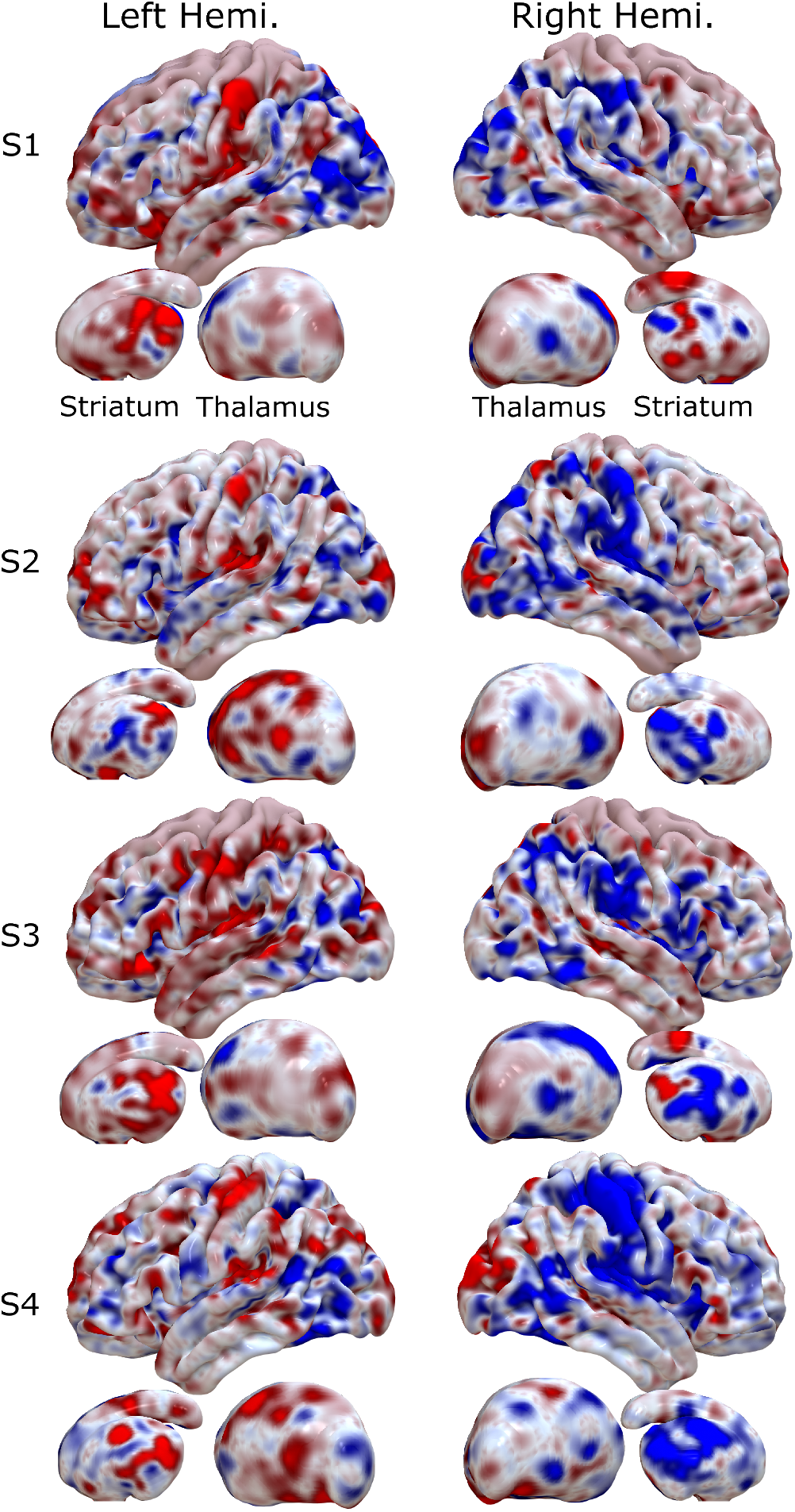
Encoding maps in standardized space for each participant. Rows represent individual participants. Columns refer to left and right views of the whole brain. Thalamus and striatum are shown beneath each cortical map. Values are z-scored.

**Supp. Fig. 5.**
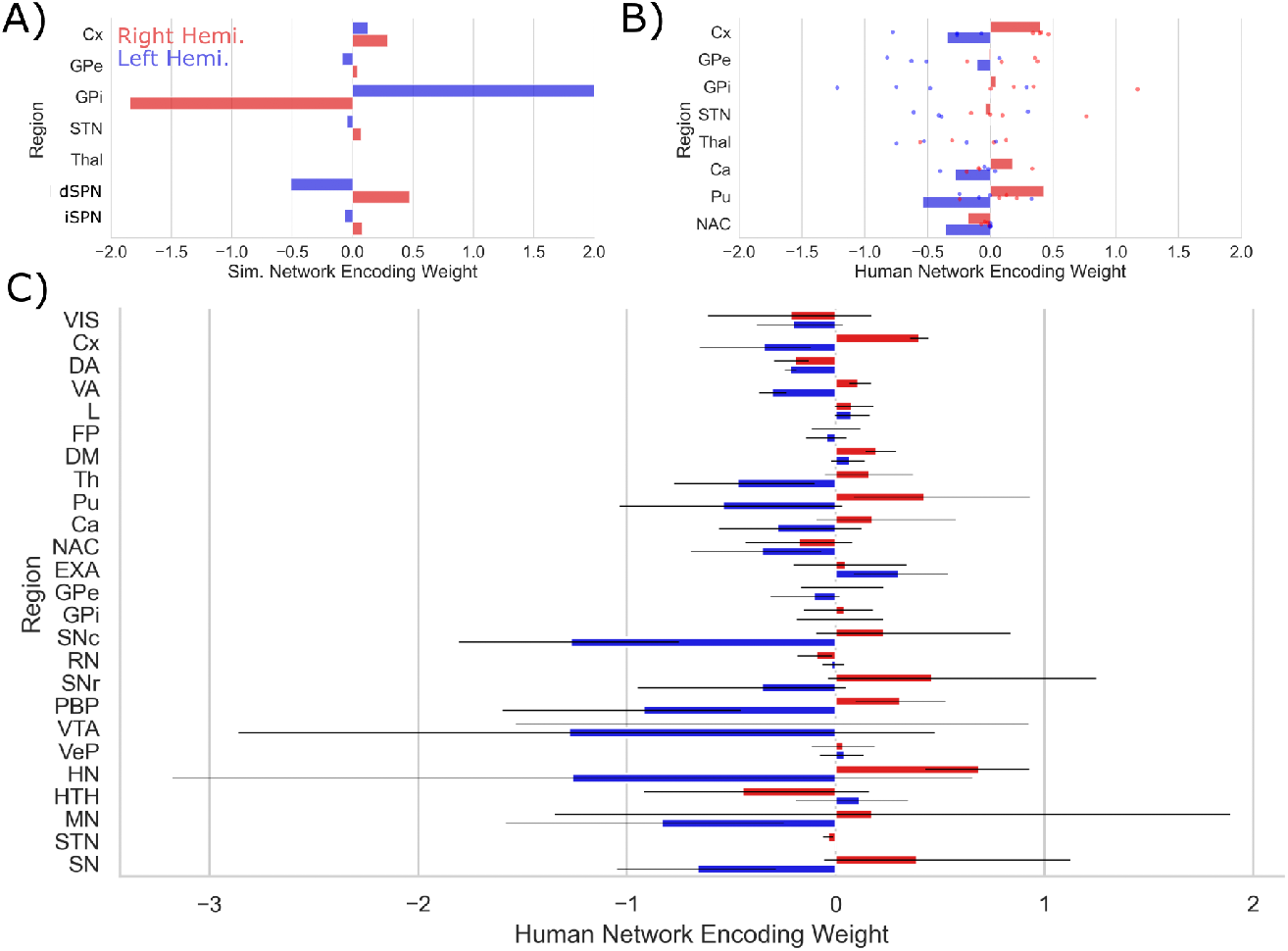
Encoding patterns by CBGT node. A) Simulated CBGT encoding weights. B) Human CBGT encoding weights for comparison with the simulated CBGT network results. Each point represents the average result for each participant. Bars represent participant-averaged data. C) The full set of human CBGT encoding weights for all captured nodes from whole-brain imaging. Gray error bars represent 95% CIs over participants. Left hemisphere weights are marked in blue and right hemisphere weights are marked in red.

### Tables

**Supp. Table 1.** Behavior. Simulated and human reaction time (in seconds) and accuracy.

| Simulated |  |  | Human |  |  |
| --- | --- | --- | --- | --- | --- |
| Sim. Part. | RT(s) | Accuracy | Human Part. | RT (s) | Accuracy |
| 1 | 0.604 | 0.59 | 1 | 0.553 | 0.538 |
| 2 | 0.559 | 0.624 | 2 | 0.537 | 0.541 |
| 3 | 0.608 | 0.61 | 3 | 0.531 | 0.553 |
| 4 | 0.596 | 0.648 | 4 | 0.54 | 0.511 |
| All | 0.592 ± 0.176 | 0.618 | All | 0.540 ± 0.076 | 0.536 |

**Supp. Table 2.** Model fits. Deviance Information Criterion (DIC) values for regression models tested.

| **Simulated** |  |  |  |  |
| --- | --- | --- | --- | --- |
| | $\Delta B$ | $\Omega$ | $\Delta DIC_{\text{null}}$ | $\Delta DIC_{\text{best}}$ |
| I | v | a | $-29.85 \pm 12.76$ | $-4.49 \pm 5.91$ |
| II | a | v | $-23.94 \pm 22.56$ | $-10.40 \pm 11.22$ |
| III | - | v | $-6.16 \pm 4.24$ | $-28.19 \pm 13.62$ |
| IV | v | - | $-22.60 \pm 7.28$ | $-11.74 \pm 14.80$ |
| V | - | a | $-7.04 \pm 11.06$ | $-27.30 \pm 8.16$ |
| VI | a | - | $-17.72 \pm 21.49$ | $-16.62 \pm 11.88$ |
| VII | - | - | $0.00 \pm 0.00$ | $-34.34 \pm 15.97$ |

**Supp. Table 2. Model fits.** Deviance Information Criterion (DIC) values for regression models 440 tested.
| **Human** |  |  |  |  |
| --- | --- | --- | --- | --- |
| | $\Delta B$ | $\Omega$ | $\Delta DIC_{\text{null}}$ | $\Delta DIC_{\text{best}}$ |
| I | v | a | $-14.90 \pm 20.58$ | $-1.52 \pm 1.04$ |
| II | a | v | $-0.44 \pm 1.11$ | $-15.99 \pm 18.56$ |
| III | - | v | $-1.47 \pm 1.30$ | $-14.96 \pm 18.56$ |
| IV | v | - | $-13.80 \pm 16.61$ | $-2.63 \pm 3.62$ |
| V | - | a | $-1.03 \pm 4.46$ | $-15.40 \pm 15.60$ |
| VI | a | - | $1.00 \pm 0.71$ | $-17.42 \pm 19.52$ |
| VII | - | - | $0.00 \pm 0.00$ | $-16.43 \pm 19.53$ |

**Supp Table 3.**
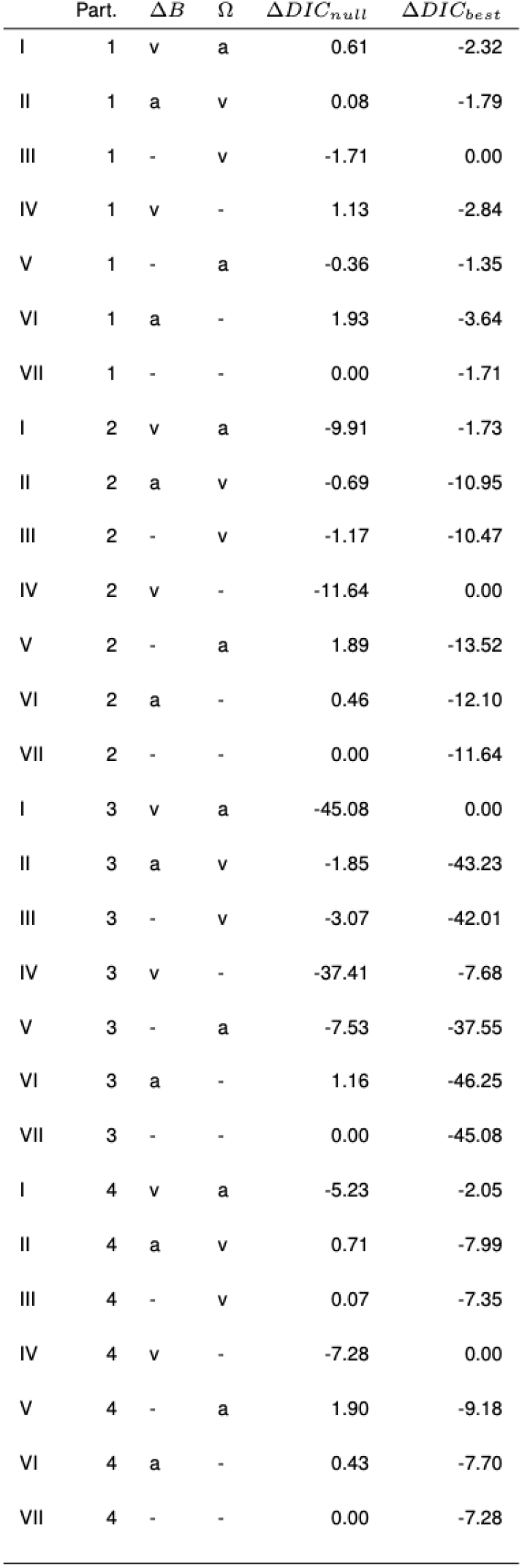
Human model fits by participant.

| | Part. | $\Delta B$ | $\Omega$ | $\Delta DIC_{null}$ | $\Delta DIC_{best}$ |
| --- | --- | --- | --- | --- | --- |
| I | 1 | v | a | 0.61 | -2.32 |
| II | 1 | a | v | 0.08 | -1.79 |
| III | 1 | - | v | -1.71 | 0.00 |
| IV | 1 | v | - | 1.13 | -2.84 |
| V | 1 | - | a | -0.36 | -1.35 |
| VI | 1 | a | - | 1.93 | -3.64 |
| VII | 1 | - | - | 0.00 | -1.71 |
| I | 2 | v | a | -9.91 | -1.73 |
| II | 2 | a | v | -0.69 | -10.95 |
| III | 2 | - | v | -1.17 | -10.47 |
| IV | 2 | v | - | -11.64 | 0.00 |
| V | 2 | - | a | 1.89 | -13.52 |
| VI | 2 | a | - | 0.46 | -12.10 |
| VII | 2 | - | - | 0.00 | -11.64 |
| I | 3 | v | a | -45.08 | 0.00 |
| II | 3 | a | v | -1.85 | -43.23 |
| III | 3 | - | v | -3.07 | -42.01 |
| IV | 3 | v | - | -37.41 | -7.68 |
| V | 3 | - | a | -7.53 | -37.55 |
| VI | 3 | a | - | 1.16 | -46.25 |
| VII | 3 | - | - | 0.00 | -45.08 |
| I | 4 | v | a | -5.23 | -2.05 |
| II | 4 | a | v | 0.71 | -7.99 |
| III | 4 | - | v | 0.07 | -7.35 |
| IV | 4 | v | - | -7.28 | 0.00 |
| V | 4 | - | a | 1.90 | -9.18 |
| VI | 4 | a | - | 0.43 | -7.70 |
| VII | 4 | - | - | 0.00 | -7.28 |

## Materials & Methods

### Simulations

We simulated neural dynamics and behavior using a biologically based, spiking cortico-basal ganglia-thalamic (CBGT) network model (*11, 19*). The network representing the CBGT circuit is composed of 9 neural populations: cortical interneurons (CxI), excitatory cortical neurons (Cx), striatal D1/D2-spiny projection neurons (dSPNs/iSPNs), striatal fast-spiking interneurons (FSI), the internal (GPi) and external globus pallidus (GPe), the subthalamic nucleus (STN), and the thalamus (Th). All the neuronal populations are segregated into two action channels with the exception of cortical (CxI) and striatal interneurons (FSIs). Each neuron in the population was modeled with an integrate-fire-or-burst-model (*50*), and a conductance-based synapse model was used for NMDA, AMPA and GABA receptors. The neuronal and network parameters (inter-nuclei connectivity and synaptic strengths) were tuned to obtain realistic baseline firing rates for all the nuclei. The details of the model are described in our previous work (*19*) as well as in the appendix for the sake of completeness.

Corticostriatal weights for D1 and D2 neurons in striatum were modulated by phasic dopamine to model the influence of reinforcement learning on network dynamics. The details of STDP learning are described in detail in previous work (*38*), but key details are shown below. As a result of these features of the CBGT network, it was capable of learning under realistic experimental paradigms with probabilistic reinforcement schemes (i.e. under reward probabilities and unstable action-outcome values).

### Threshold for CBGT network decisions

A decision between the two competing actions (“left” and “right”) was considered to be made when either of the thalamic subpopulations reached a threshold of 30Hz. This threshold was set based on the network dynamics for the chosen parameters with a aim to obtain realistic reaction times. The maximum time allowed to reach a decision was 1000ms. If none of the thalamic subpopulations reach the threshold of 30Hz, no action was considered to be taken. Such trials were dropped from further analysis. Reaction/decision times were calculated as time from stimulus onset to decision (either subpopulation reaches the threshold). The “slow” and “fast” trials were categorized as reaction times ≥ 75th percentile (314.5ms) and reactions time *<* 50th percentile (196.0ms), respectively, of the reaction time distributions. The firing rates of the CBGT nuclei during the reaction times were used for prediction analysis as discussed in Section.

### Corticostriatal weight plasticity

The corticostriatal weights are modified by a dopamine-mediated STDP rule, where the phasic dopamine is modulated by reward prediction error. The internal estimate of the reward is calculated at every trial by a Q-learning algorithm which is subtracted from the reward associated with the experimental paradigm to yield a trial-by-trial estimate of the reward prediction error. The effect of dopaminergic release is receptor dependent; a rise in dopamine promotes potentiation for D1-SPNs and depression for D2-SPNs. The degree of change in the weights is dependent on an eligibility trace which is proportional to the coincidental pre-synaptic (cortical) and post-synaptic (striatal) firing rates. The STDP rule is described in detail in (*38*) as well as in the appendix.

### In silico experimental design

We follow the paradigm of a 2 arm bandit task, where the CBGT network learns to consistently choose the rewarded action until the block changes (i.e the reward contingencies switch), at which point the CBGT network re-learns the rewarded action (reversal learning). Each session consists of 40 trials with a block change every 10 trials. The reward probabilities represent a conflict of (75%, 25%); that is, in a left block, 75% of the left actions are rewarded, whereas 25% of the right actions are rewarded. The inter-trial-interval in network time is fixed to 600ms.

To maximize the similarity between the CBGT network simulations and our human data, we randomly varied the initialization of the network such that neurons with a specific connection probability were randomly chosen for each simulated subject, with the background input to the nuclei for each simulated subject as a mean-reverting random walk (noise was drawn from the normal distribution N(0,1)). These means are listed in Supp. Table 4.

### Participants

Four neurologically healthy adult human primates (two female, all right-handed, 29-34 years old) were recruited and paid $30 per session, in addition to a performance bonus and a bonus for completing all nine sessions. These participants were recruited from the local university population.

All procedures were approved by the Carnegie Mellon University Institutional Review Board. All research participants provided informed consent to participate in the study and consent to publish any research findings based on their provided data.

### Experimental design

The experiment used male and female Greebles (*44*) as selection targets. Participants were first trained to discriminate between male and female Greebles to prevent errors in perceptual discrimination from interfering with selection on the basis of value. Using a two-alternative forced choice task, participants were presented with a male and female Greeble and asked to select the female, with the male and female Greeble identities resampled on each trial. Participants received binary feedback regarding their selection (correct or incorrect). This criterion task ended after participants reached 95% accuracy. After reaching perceptual discrimination criterion for each session, each participant was tested under nine reinforcement learning conditions composed of 300 trials each, generating 2700 trials per participant in total. Data were collected from four participants in accordance with a replication-based design, with each participant serving as a replication experiment. Participants completed these sessions in randomized order. Each learning trial presented a male and female Greeble (*44*), with the goal of selecting the gender identity of the Greeble that was most rewarding. Because individual Greeble identities were resampled on each trial, the task of the participant was to choose the gender identity rather than the individual identity of the Greeble which was most rewarding.

Probabilistic reward feedback was given in the form of points drawn from the normal distribution *𝒩* (*μ* = 3, *σ* = 1) and converted to an integer. These points were displayed at the center of the screen. For each run, participants began with 60 points and lost one point for each incorrect decision. To promote incentive compatibility (*51, 52*), participants earned a cent for every point earned. Reaction time was constrained such that participants were required to respond within between 0.1 s and 0.75 s from stimulus presentation. If participants responded in ≤ 0.1 s, ≥ 0.75 s, or failed to respond altogether, the point total turned red and decreased by 5 points. Each trial lasted 1.5 s and reward feedback for a given trial was displayed from the time of the participant’s response to the end of the trial. To manipulate change point probability, the gender identity of the most rewarding Greeble was switched probabilistically, with a change occurring every 10, 20, or 30 trials, on average. To manipulate the belief in the value of the optimal target, the probability of reward for the optimal target was manipulated, with *P* set to 0.65, 0.75, or 0.85. Each session combined one value of *P* with one level of volatility, such that all combinations of change point frequency and reward probability were imposed across the nine sessions. Finally, the position of the high-value target was pseudo-randomized on each trial to prevent prepotent response selections on the basis of location.

### Behavioral analysis

Statistical analyses and data visualization were conducted using custom scripts written in R (R Foundation for Statistical Computing, version 3.4.3) and Python (Python Software Foundation, version 3.5.5). Binary accuracy data were submitted to a mixed effects logistic regression analysis with either the degree of conflict (the probability of reward for the optimal target) or the degree of volatility (mean change point frequency) as predictors. The resulting log-likelihood estimates were transformed to likelihood for interpretability. RT data were log-transformed and submitted to a mixed effects linear regression analysis with the same predictors as in the previous analysis. To determine if participants used ideal observer estimates to update their behavior, two more mixed effects regression analyses were performed. Estimates of change point probability and the belief in the value of the optimal target served as predictors of reaction time and accuracy across groups. As before, we used a mixed logistic regression for accuracy data and a mixed linear regression for reaction time data.

### Estimating evidence accumulation using drift diffusion modeling

To assess whether and how much the ideal observer estimates of change point probability (Ω) and the belief in the value of the optimal target (ΔB) (*3, 7*) updated the rate of evidence accumulation (*v*), we regressed the change-point-evoked ideal observer estimates onto the decision parameters using hierarchical drift diffusion model (HDDM) regression (*53*). These ideal observer estimates of environmental uncertainty served as a more direct and continuous measure of the uncertainty we sought to induce with our experimental manipulations. Using this more direct approach, we pooled change point probability and belief across all conditions and used these values as our predictors of drift rate and boundary height. Responses were accuracy-coded, and the belief in the difference between targets values was transformed to the belief in the value of the optimal target (Δ*B*_optimal(t)_ = *B*_optimal(t)_ − *B*_suboptimal(t)_). This approach allowed us to estimate trial-by-trial covariation between the ideal observer estimates and the decision parameters.

To find the models that best fit the observed data, we conducted a model selection process using Deviance Information Criterion (DIC) scores. A lower DIC score indicates a model that loses less information. Here, a difference of two points from the lowest-scoring model cannot rule out the higher scoring model; a difference of three to seven points suggests that the higher scoring model has considerably less support; and a difference of 10 points suggests essentially no support for the higher scoring model (*43, 54*). We evaluated the DIC scores for the set of fitted models relative to an intercept-only regression model 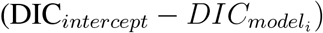.

### MRI Data Acquisition

Neurologically healthy human participants (N=4, 2 female) were recruited. Each participant was tested in nine separate imaging sessions using a 3T Siemens Prisma scanner. Session 1 included a set of anatomical and functional localizer sequences (e.g., visual presentation of Greeble stimuli with no manual responses, and left vs. right button responses to identify motor networks). Sessions 2-10 collected five functional runs of the dynamic 2-armed bandit task (60 trials per run). Male and female “greebles” served as the visual stimuli for the selection targets (*44*), with each presented on one side of a central fixation cross. Participants were trained to respond within 1.5 seconds.

To minimize the convolution of the hemodynamic response from trial to trial, inter-trial intervals were sampled according to a truncated exponential distribution with a minimum of 4 s between trials, a maximum of 16 s, and a rate parameter of 2.8 s. To ensure that head position was stabilized and stable over sessions, a CaseForge head case was customized and printed for each participant. The task-evoked hemodynamic response was measured using a high spatial (2*mm*^3^ voxels) and high temporal (750ms TR) resolution echo planar imaging approach. This design maximized recovery of single-trial evoked BOLD responses in subcortical areas, as well as cortical areas with higher signal-to-noise ratios. During each functional run, eye-tracking (EyeLink, SR Research Inc.), physiological signals (ECG, respiration, and pulse-oximetry via the Siemens PMU system) were also collected for tracking attention and for artifact removal.

### Preprocessing

fMRI data were preprocessed using the default pipeline of fMRIPrep (*55*), a standard toolbox for fMRI data preprocessing that provides stability to variations in scan acquisition protocols, a minimal user manipulation, and easily interpretable, comprehensive output results reporting.

### Single-trial response estimation

By means of a univariate general linear model (GLM) within participant trial-wise responses at the voxel-level were estimated. Specifically, for each fMRI run preprocessed BOLD time series were regressed onto a design matrix, where each task trial corresponded to a different column, and was modeled using a boxcar function convolved with the default hemodynamic response function given in SPM12. Thus, each column in the design matrix estimated the average BOLD activity within each trial. In order to account for head motion, the six realignment parameters (3 rotations, 3 translations) were included as covariates. In addition, a high-pass filter (128 s) was applied to remove low-frequency artifacts. Parameter and error variance were estimated using the RobustWLS toolbox, which adjusts for further artifacts in the data by inversely weighting each observation according to its spatial noise (*56*).

Finally, estimated trial-wise responses were concatenated across runs and sessions and then stacked across voxels to give a matrix, 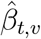, of T (trial estimations) x V (voxels) for each participant.

### Single-trial response prediction

A machine learning approach was applied to predict left/right greeble choices from the trialwise responses. First, using the trial-wise hemodynamic responses, we estimated the contrast in neural activation when the participant made a left versus right selection. A Lasso-PCR classifier (i.e. an L1-constrained principal component logistic regression) was estimated for each participant according to the below procedure.

First, a singular value decomposition (SVD) was applied to the input matrix *X*:

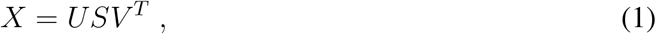

where the product matrix *Z* = *US* represents the principal component scores, i.e. the projected values of *X* into the principal component space, and *V*^*T*^ an orthogonal matrix whose rows are the principal directions in feature space. Then the binary response variable *y* (Left/Right choice) was regressed onto *Z*, where the estimation of the *β* coefficients is participant to a L1 penalty term C in the objective function:

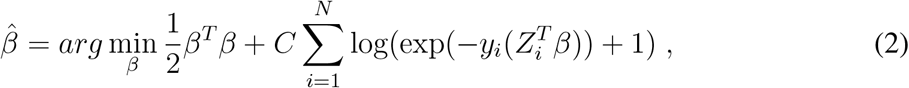

where *β* and Z include the intercept term, *y*_*i*_ = {−1, 1} and N is the number of observations. Projection of the estimated 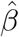 coefficients back to the original feature (voxel) space was done to yield a weight map 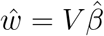, which in turn was used to generate final predictions *ŷ*:

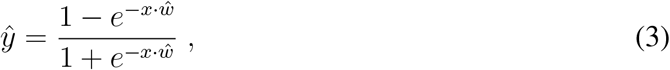

where *x* denotes the vector of voxel-wise responses for a given trial (i.e. a given row in the *X* matrix). When visualizing the resulting weight maps, these were further transformed to encoded brain patterns. This step was performed to aid in correct interpretation in terms of the studied brain process, because doing this directly from the observed weights in multivariate classification (and regression) models can be problematic (*57*).

Here, the competition between left-right neural responses decreases classifier decoding accuracy, as neural activation associated with these actions becomes less separable. Therefore, classifier prediction serves as a proxy for response competition. To quantify uncertainty from this, we calculated the Euclidean distance of these decoded responses *ŷ* from the statistically optimal choice on a given trial, *opt choice*. This yielded a trial-wise uncertainty metric derived from the decoded competition between neural responses.

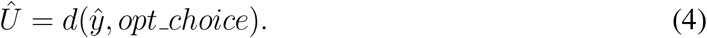

The same analytical pipeline was used to calculate single trial responses for simulated data with a difference that trial-wise average firing rates of all nuclei from the simulations were used instead of fMRI hemodynamic responses.

### Neuron model

We used integrate-and-fire-or-burst model that models the membrane potential *V* (*t*) as

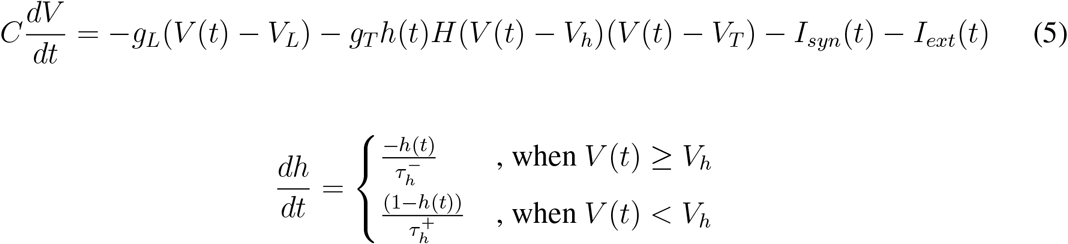

where *g*_*L*_ represents the leak conductance, *V*_*L*_ is the leak reversal potential and the first term *g*_*L*_(*V* (*t*) − *V*_*L*_) is the leak current; a low threshold *Ca*^2+^ current with maximum conductance as *g*_*T*_, gating variable *h*(*t*), a heaviside function *H*, reversal potential *V*_*T*_ ; *I*_*syn*_ is the synaptic current and *I*_*ext*_ is the external current. This neuron model is capable of producing post inhibitory bursts, regulated by the gating variable that decays with the time constant 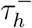, when the membrane potential reaches a certain threshold *V*_*h*_ and rises with time constant 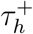. However, when *g*_*T*_ is set to zero, the neuronal dynamics reduce to a leaky integrate and fire neuron. Currently, we model GPe and STN neuronal populations with bursty neurons and the remaining neuronal populations with leaky integrate-and-fire neurons, with conductance-based synapses.

The synaptic current *I*_*syn*_(*t*) consists of three components, two excitatory currents corresponding to AMPA and NMDA receptors and one inhibitory current corresponding to GABA receptors, and is calculated as below:

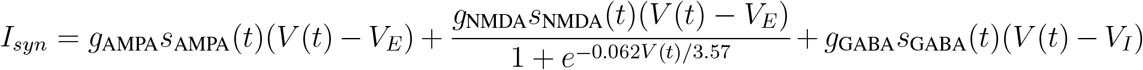

where *g*_i_ represents the maximum conductance corresponding to the receptor *i* ∈ (AMPA, NMDA and GABA), *V*_*I*_ and *V*_*E*_ represent the excitatory and inhibitory reversal potentials, and *s*_i_ represents the gating variable for the channels, with dynamics given by:

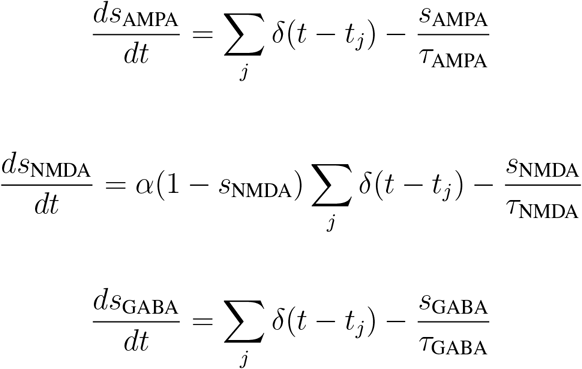

The gating variables for AMPA and GABA acts as leaky integrators that are increased by all incoming spikes, with an additional constraint for NMDA that ensures that the maximum value of *s*_NMDA_ remains below 1.

The values of neuronal parameters for all the nuclei are listed in Table S4, external inputs to the CBGT nuclei are listed in Table S5, and the synaptic parameter values are listed in Table S6.

**Supp. Table 4.** Neuronal parameters. Neuronal parameters for each nucleus are listed in the left column, with values shown on the right.

| Parameter | unit | Cx | Cxl | dSPN | iSPN | FSI | GPe | STN | Thalamus |
| --- | --- | --- | --- | --- | --- | --- | --- | --- | --- |
| $\tau_m$ (membrane time constant) | ms | 20 | 10 | 20 | 20 | 10 | 20 | 20 | 27.78 |
| $V_{rest}$ (resting membrane potential) | mV | -70 | -70 | -70 | -70 | -70 | -70 | -70 | -70 |
| $V_{threshold}$ (threshold potential) | mV | -50 | -50 | -50 | -50 | -50 | -50 | -50 | -50 |
| $V_L$ (leak reversal) | mV | -55 | -55 | -55 | -55 | -55 | -55 | -55 | -55 |
| $g_T$ (low threshold $Ca^{2+}$ maximal conductance) | mS/cm<sup>2</sup> | 0 | 0 | 0 | 0 | 0 | 0.06 | 0.06 | 0 |
| $V_h$ (threshold potential for burst activation) | mV | -60 | -60 | -60 | -60 | -60 | -60 | -60 | -60 |
| $V_T$ ($Ca^{2+}$ reversal potential) | mV | 120 | 120 | 120 | 120 | 120 | 120 | 120 | 120 |
| $\tau_h^-$ (burst duration in ms) | ms | 20 | 20 | 20 | 20 | 20 | 20 | 20 | 20 |
| $\tau_h^+$ (hyperpolarization duration) | ms | 100 | 100 | 100 | 100 | 100 | 100 | 100 | 100 |

### Spike timing dependent plasticity rule

The plasticity rule we use is a dopamine modulated STDP rule also described in (*38*). All the values of the relevant parameters are listed in Table S8. The weight update of a corticostriatal synapse is controlled by three factors: 1) an eligibility trace, 2) the type of the striatal neuron (iSPN/dSPN), and 3) the level of dopamine.

To compute these quantities for a given synapse, an activity trace of each neuron in the pre-synaptic and post-synaptic populations is tracked via the equations

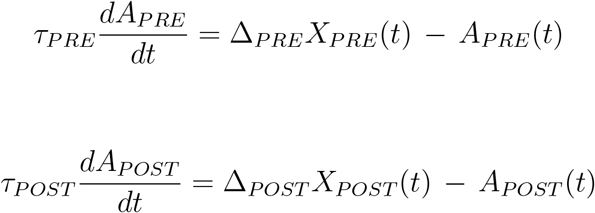

where *X*_*PRE*_, *X*_*POST*_ are spike trains, such that *A*_*PRE*_ and *A*_*POST*_ maintain a filtered record of synaptic spiking of the pre/post neuron, respectively, with spike impact parameters Δ_*PRE*_, Δ_*POST*_ and time constants *τ*_*PRE*_, *τ*_*POST*_.

If the post-synaptic spike follows the spiking activity of the pre-synaptic population closely enough in time, then eligibility trace (*E*) increases and allows for plasticity to occur. On the other hand, if a pre-synaptic spike follows the spiking activity of the post-synaptic population, then *E* decreases. In absence of any activity and spikes, the eligibility trace decays to zero with a time constant *τ*_*E*_. Putting these effets together, we obtain the equation

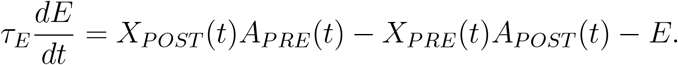

The synaptic weight update depends on the dopamine receptor type of the striatal neuron; that is, if the neuron is a dSPN or iSPN. We assume that a phasic dopamine release promotes long term potentiation (LTP) in dSPNs and long term depression (LTD) in iSPNs. This factor is indicated by the learning rate parameter *α*_*w*_, which is set to a positive value for dSPNs and a negative value for iSPNs. The weight update dynamics is given by:

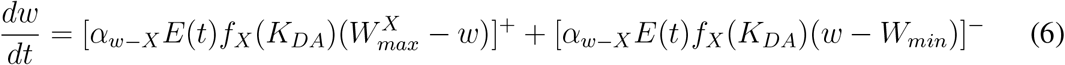

where *X* ∈ { dSPN, iSPN } with *α*_*w*–*dSPN*_ > 0 and *α*_*w*–*iSPN*_ < 0. Here, the weights of the corticostriatal synapses are bounded between the maximal value 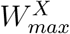, which depends on the SPN type, and a minimal value of *W*_*min*_ = 0.001. The precise values used for all relevant parameters are listed in Table S8.

In the weight update rule (6), *K*_*DA*_ represents the dopamine level present. This quantity changes as a result of phasic release of dopamine (increments of size *DA*_*inc*_), which is correlated to the reward prediction error encountered in the environment. The parameter *C*_*scale*_ defines the scaling between the reward prediction error and the amount of dopamine released, and *K*_*DA*_ obeys the equation

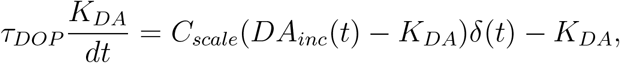

where

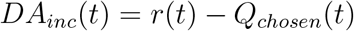

for reward *r*(*t*) and expected value *Q*_*chosen*_(*t*) of the chosen action. Trial-by-trial estimates of the values of the actions (left/right) are maintained by a simple Q-update rule:

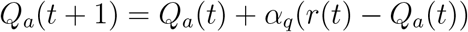

where *a* ∈ {left, right} and where *α*_*q*_ represents the learning rate of the Q-values.

Finally, the function *f*_*X*_(*K*_*DA*_) converts the level of dopamine into an impact on plasticity in a way that depends on the identity *X* of the post-synaptic neuron, as follows:

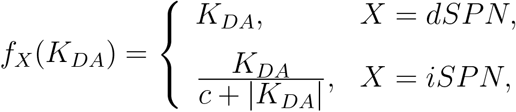

where *c* sets the dopamine level where *f*_*iSPN*_ reaches half-maximum.

**Supp. Table 5.**
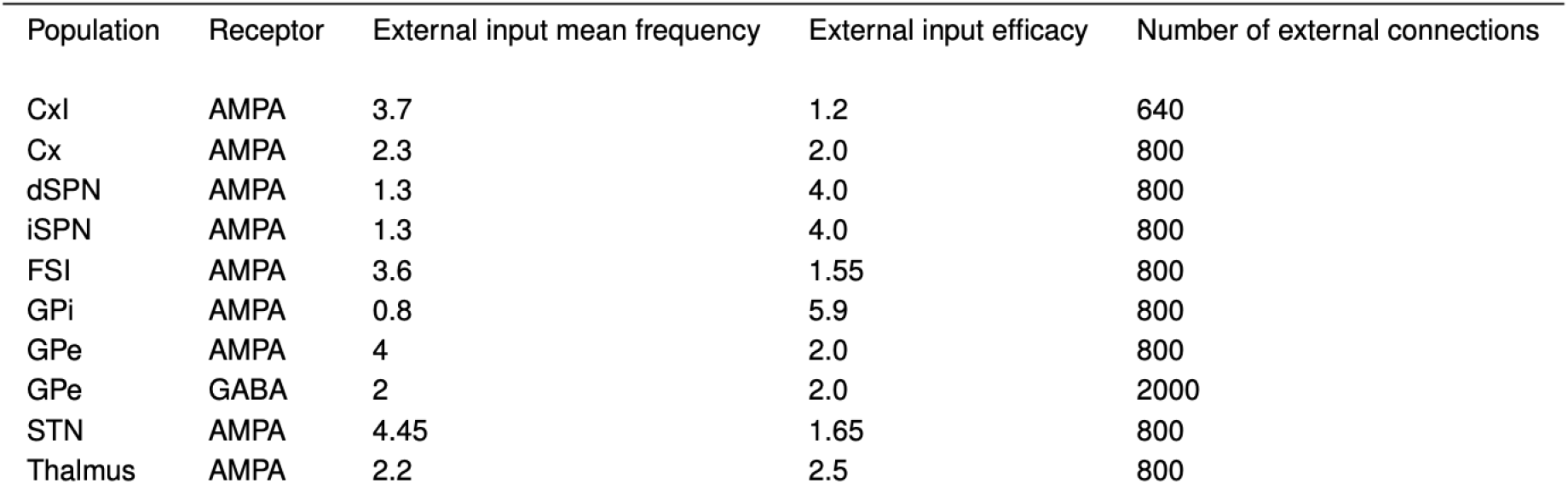
External inputs to CBGT nuclei. Each nucleus is listed on the left, with input frequency, efficacy, and number of connections listed by receptor.

**Supp. Table 6.** Synaptic parameters. Parameters for the simulated synapses.

| Parameter | unit | Value |
| --- | --- | --- |
| $\tau_{\text{AMPA}}$ | ms | 2 |
| $V_E$ | mV | 0 |
| $\tau_{\text{NMDA}}$ | ms | 100 |
| $\tau_{\text{GABA}}$ | ms | 5 |
| $V_I$ | mV | -70 |
| $\alpha$ | - | 0.6332 |

**Supp. Table 7.** CBGT connectivity. Connection type and probability by nucleus and receptor.

| Connection type | Connection probability | g (nS) | Receptor |
| --- | --- | --- | --- |
| Cx-dSPN | 1.0 | 0.015 | AMPA |
| Cx-dSPN | 1.0 | 0.02 | NMDA |
| Cx-iSPN | 1.0 | 0.015 | AMPA |
| Cx-iSPN | 1.0 | 0.02 | NMDA |
| Cx-FSI | 1.0 | 0.43 | AMPA |
| Cx-Th | 1.0 | 0.025 | AMPA |
| Cx-Th | 1.0 | 0.035 | NMDA |
| Cx-Cx | 0.13 | 0.0127 | AMPA |
| Cx-Cx | 0.13 | 0.08 | NMDA |
| Cx-CxI | 0.0725 | 0.113 | AMPA |
| Cx-CxI | 0.0725 | 0.525 | NMDA |
| CxI-Cx | 0.5 | 1.05 | GABA |
| CxI-CxI | 1.0 | 1.075 | GABA |
| dSPN-dSPN | 0.45 | 0.28 | GABA |
| dSPN-iSPN | 0.45 | 0.28 | GABA |
| dSPN-GPi | 1.0 | 2.09 | GABA |
| iSPN-iSPN | 0.45 | 0.28 | GABA |
| iSPN-dSPN | 0.5 | 0.28 | GABA |
| iSPN-GPe | 1.0 | 4.07 | GABA |
| FSI-FSI | 1.0 | 3.2583 | GABA |
| FSI-dSPN | 1.0 | 1.77 | GABA |
| FSI-iSPN | 1.0 | 1.66 | GABA |
| GPe-GPe | 0.067 | 1.75 | GABA |
| GPe-STN | 0.067 | 0.35 | GABA |
| GPe-GPi | 1.0 | 0.058 | GABA |
| STN-GPe | 0.1617 | 0.07 | AMPA |
| STN-GPe | 0.1617 | 1.51 | NMDA |
| STN-GPi | 1.0 | 0.038 | GABA |
| GPi-Th | 1.0 | 0.033 | GABA |
| Th-dSPN | 1.0 | 0.38 | AMPA |
| Th-iSPN | 1.0 | 0.38 | AMPA |
| Th-FSI | 0.83 | 0.1 | AMPA |
| Th-Cx | 0.83 | 0.03 | NMDA |

**Supp. Table 8.** Number of neurons in each CBGT nucleus.

| Population | Number of neurons |
| --- | --- |
| Cx | 204 |
| CxI | 186 |
| dSPN | 75 |
| iSPN | 75 |
| FSI | 75 |
| GPe | 750 |
| GPI | 75 |
| STN | 750 |
| Th | 75 |

**Supp. Table 9.**
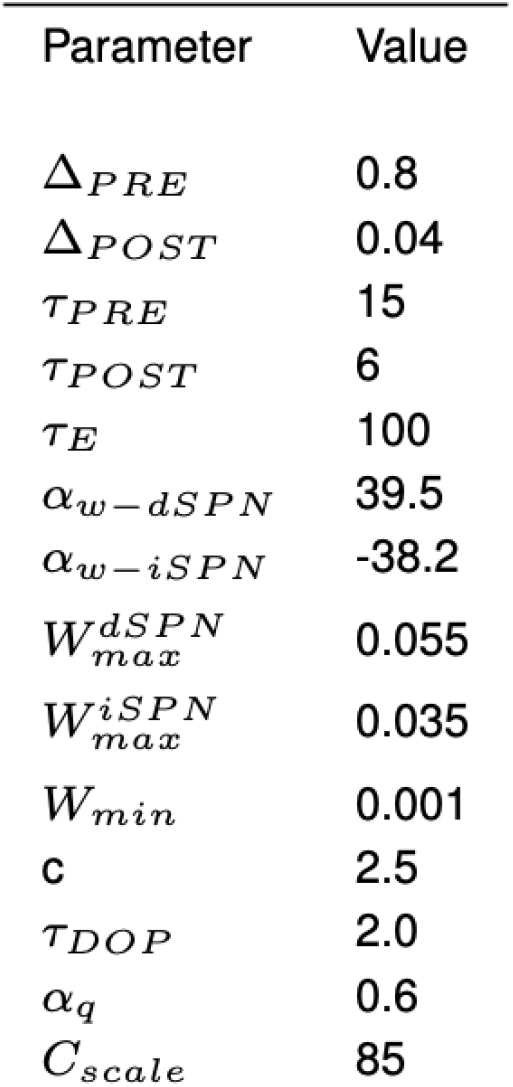
STDP parameters.

| Parameter | Value |
| --- | --- |
| $\Delta_{PRE}$ | 0.8 |
| $\Delta_{POST}$ | 0.04 |
| $\tau_{PRE}$ | 15 |
| $\tau_{POST}$ | 6 |
| $\tau_E$ | 100 |
| $\alpha_{w-dSPN}$ | 39.5 |
| $\alpha_{w-iSPN}$ | -38.2 |
| $W_{max}^{dSPN}$ | 0.055 |
| $W_{max}^{iSPN}$ | 0.035 |
| $W_{min}$ | 0.001 |
| **c** | 2.5 |
| $\tau_{DOP}$ | 2.0 |
| $\alpha_q$ | 0.6 |
| $C_{scale}$ | 85 |

